# The genomic basis of adaptation to the deep water ‘twilight zone’ in Lake Malawi cichlid fishes

**DOI:** 10.1101/102830

**Authors:** Christoph Hahn, Martin J Genner, George F Turner, Domino A Joyce

**Affiliations:** Evolutionary and Environmental Genomics Group (@EvoHull), School of Environmental Sciences, University of Hull, Hull HU5 7RX, UK.; Institute of Zoology, University of Graz, A-8010 Graz, Austria.; School of Biological Sciences, University of Bristol, Bristol Life Sciences Building, 24 Tyndall Avenue, Bristol. BS8 1TQ. UK.; School of Biological Sciences, Bangor University, Bangor, Gwynedd LL57 2UW, Wales, UK.

**Keywords:** Root effect, supergene, cichlid, hemoglobin, haemoglobin, sensory drive

## Abstract

Deep water environments are characterized by low levels of available light at increasingly narrow spectra, great hydrostatic pressure and reduced dissolved oxygen - conditions predicted to exert highly specific selection pressures. In Lake Malawi over 800 cichlid species have evolved, and this adaptive radiation extends into the “twilight zone” below 100 metres. We use population-level RAD-seq data to investigate whether four endemic deep water species (*Diplotaxodon* spp.) have experienced divergent selection within this environment. We identify candidate genes including regulators of photoreceptor function, photopigments, lens morphology and haemoglobin, many not previously implicated in cichlid adaptive radiations. Co-localization of functionally linked genes suggests co-adapted “supergene” complexes. Comparisons of *Diplotaxodon* to the broader Lake Malawi radiation using genome resequencing data revealed functional substitutions in candidate genes. Our data provide unique insights into genomic adaptation to life at depth, and suggest genome-level specialisation for deep water habitat as an important process in cichlid radiation.

---

Deep water environments pose an array of physiological challenges to the organisms that inhabit them. Below 50m, increased hydrostatic pressure, reduced levels of dissolved oxygen and a lack of ambient light, will all produce characteristic selection pressures ^1,2^. Evolutionary adaptation of species to this range of ecological challenges should be possible to detect at the genomic level, and yet surprisingly few studies have addressed this. In the context of ecological speciation and adaptive radiation, divergence along depth gradients is associated with the evolution of reproductive isolation in many marine ^3–6^ and freshwater species groups. Specifically, fish species within adaptive radiations of the freshwater Lakes Baikal, Tanganyika, Malawi differ extensively in the depth ranges that they occupy. Thus, investigating the genomic regions involved should provide powerful insights into the rapid ecological adaptation in these species, including physiological characteristics that have been subject to divergent selection.

The increase in hydrostatic pressure associated with depth affects multiple biological processes, from the activity of macromolecular protein assemblages such as tubulin and actin, to cellular processes such as osmoregulation and actin potential transmission in nervous cells ^7,8^. As a consequence, pressure can affect the nervous system, cardiac function and membrane transport systems relatively quickly, whereas other systems are more resilient to change ^9^. Coupled with the increase in pressure is a decline in the spectral range and intensity of ambient light ^10^. In aquatic systems light intensity decreases exponentially with water depth ^11^ and in clear water the long wavelength portion of the visible light spectrum (red light) is increasingly attenuated, shifting the spectral median towards short wavelength, monochromatic (blue) ‘twilight’ conditions ^12^ in deep water environments. In marine mesopelagic fishes perhaps the most recognised morphological adaptation of the visual system for life at increased depth is eye enlargement, which accommodates the reduced light intensity by increasing the chance of photon capture ^13,14^. In addition, shifts in the relative abundance of rods and cones in the retina have been associated with habitat depth ^15,16^; in vertebrates, cones mediate photopic vision under bright light conditions, while rods contain specialized pigments for scotopic vision under dim-light conditions ^17^.

Comparative analyses of the light absorption spectra of photopigments in different species have firmly established the role of “spectral tuning” in sensory adaptations, i.e. shifts in the maximum spectral sensitivity of photopigments towards the peak wavelength of the available light ^18–21^. Differential expression and alterations in amino acid sequences of opsin genes have been identified as the underlying molecular mechanisms for spectral shifts ^19,21–23^. In the context of deep water adaptations, a range of genomic modifications affecting rhodopsin genes (*Rh1* and *Rh2*) which code for the rod pigments, have been implied in spectral tuning in marine mesopelagic ^6,18,19^, and freshwater fishes, including the Lake Baikal sculpins (genus *Cottus*) ^24^ and deep-water cichlids of the East African great lakes ^25^. In this study we consider the adaptations of Lake Malawi’s twilight zone (50-200m) cichlids in more detail. This region is inhabited by members of an endemic deep-water haplochromine cichlid lineage, that includes approximately 20 species of *Diplotaxodon*, plus the closely related *Pallidochromis tokolosh,* all of which are zooplanktivorous or piscivorous ^26^. Sympatric species in the lineage often differ in male monochromatic nuptial colour and morphological traits including eye size ^27^ and this divergence must have happened since Lake Malawi achieved deep water conditions, in the last 5 million years or even more recently given evidence that the lake has been dry or shallow for much of its history ^28,29^. Currently, little empirical data on species specific depth distributions of *Diplotaxodon* are available, but there are indications that some species may breed in different depth zones ^30^. Additionally, survey catch records indicate that members of the *D. macrops* complex are found at depths of 150-220m during the day, while members of the *D. limnothrissa* complex occupy all oxygenated depths (ie. down to ~250m), with a peak of abundance at ~60m ^31^.

In this study we use population-level genome-wide SNP data to infer ecologically relevant physiological differences previously undetected between these species. We use genome scans and three independent candidate outlier approaches to test the prediction that regions of the genome involved in adaptation to depth (regions involved in adaptation to visual signal difference and hydrostatic pressure change) will be more divergent than average genomic divergence between the species. We characterize genomic variants underlying the observed interspecific eye morphological differences, and identify likely candidate regions for adaptation to life at depth.

## Results

### Population structure and genome-wide interspecific divergence

After stringent filtering the RAD data comprised 11,786 RAD tags that were each represented in at least 80% of individuals (minimum of 5) in each of the four populations (see Figure 1, Table S1). Three methods confirmed that these are indeed four different species: Maximum-likelihood inference based on a concatenated alignment of these RAD tags (total alignment length 1,053,675 bp) (Figure 2a), with branches separating species consistently receive high statistical support (bootstrap > 95%). *D.* ‘macrops black dorsal’, *D.* ‘limnothrissa black pelvic’, and *D.* ‘macrops ngulube’ were each reciprocally monophyletic, while *D.* ‘macrops offshore’ was paraphyletic with respect to *D.* ‘limnothrissa black pelvic’. Principal component analysis (PCA) based on 11,786 SNPs (using only one single SNP per RAD tag) confirmed strong population structure, with putative conspecific individuals grouping into distinct, non-overlapping clusters along the first two principal components (Figure S1). Discriminant analysis of principal components (DAPC) consistently assigned individuals with putative conspecifics (cluster assignment probability for all individuals 100%, k=4, Figure S2). Interspecific genome-wide divergence (see Table S2) ranged from F*_ST_*=0.05(*D*. ‘limnothrissa black pelvic’ vs. *D*. ‘macrops offshore’) to F*_ST_* = 0.09 (*D*. ‘limnothrissa black pelvic’ vs. *D*. ‘macrops ngulube’). Figure 2b illustrates genome wide patterns of F*_ST_* divergence between species.

**Figure 1.**
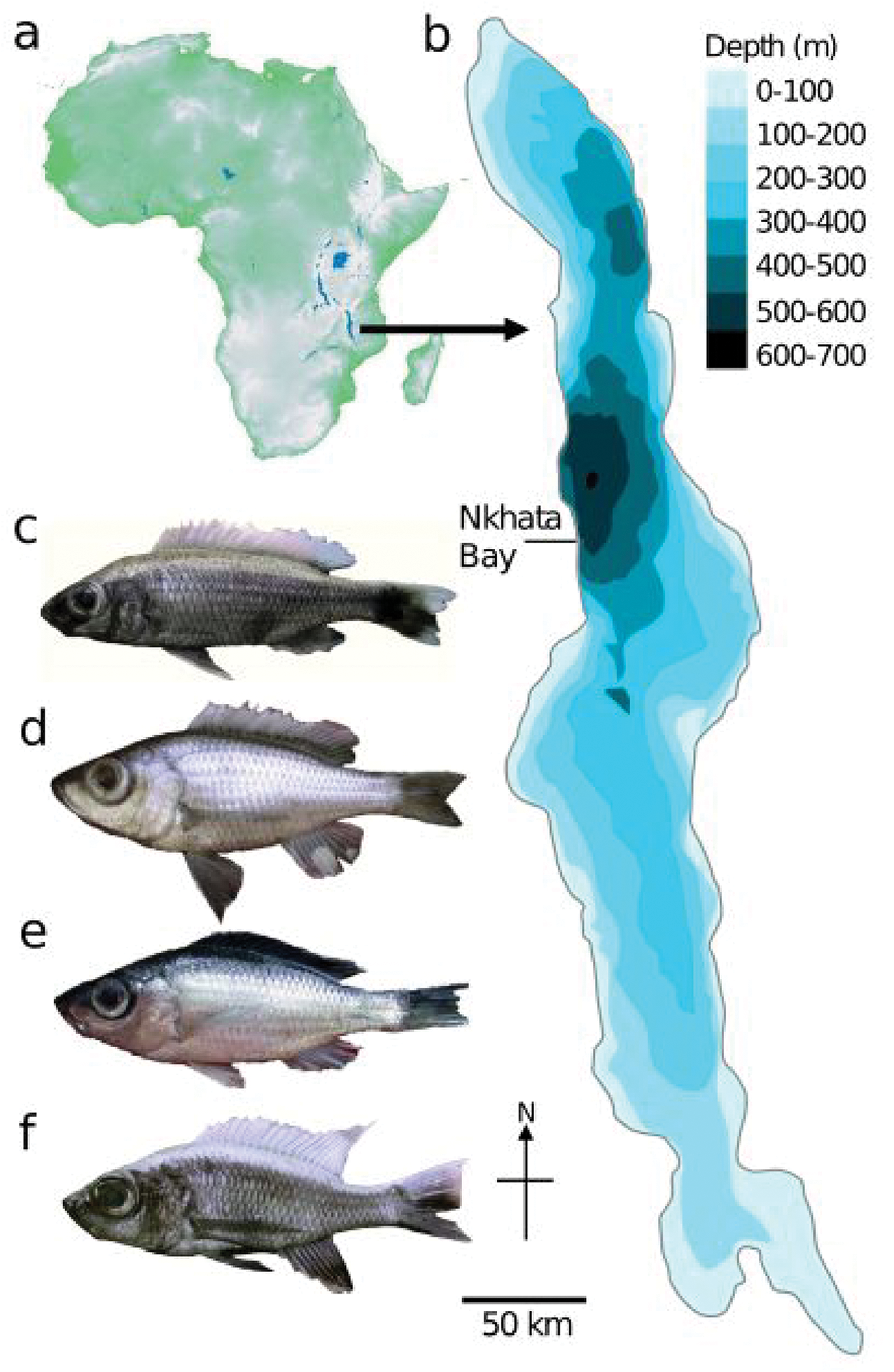
Maps of (a) Africa (topographic) and (b) Lake Malawi (bathymetric) indicating the sampling location Nkhata Bay. (c) *D*. ‘limnothrissa black pelvic’; (d) *D*. ‘macrops offshore’; (e) *D*. ‘macrops black dorsal’; (f) *D*. ‘macrops ngulube’.

**Figure 2.**
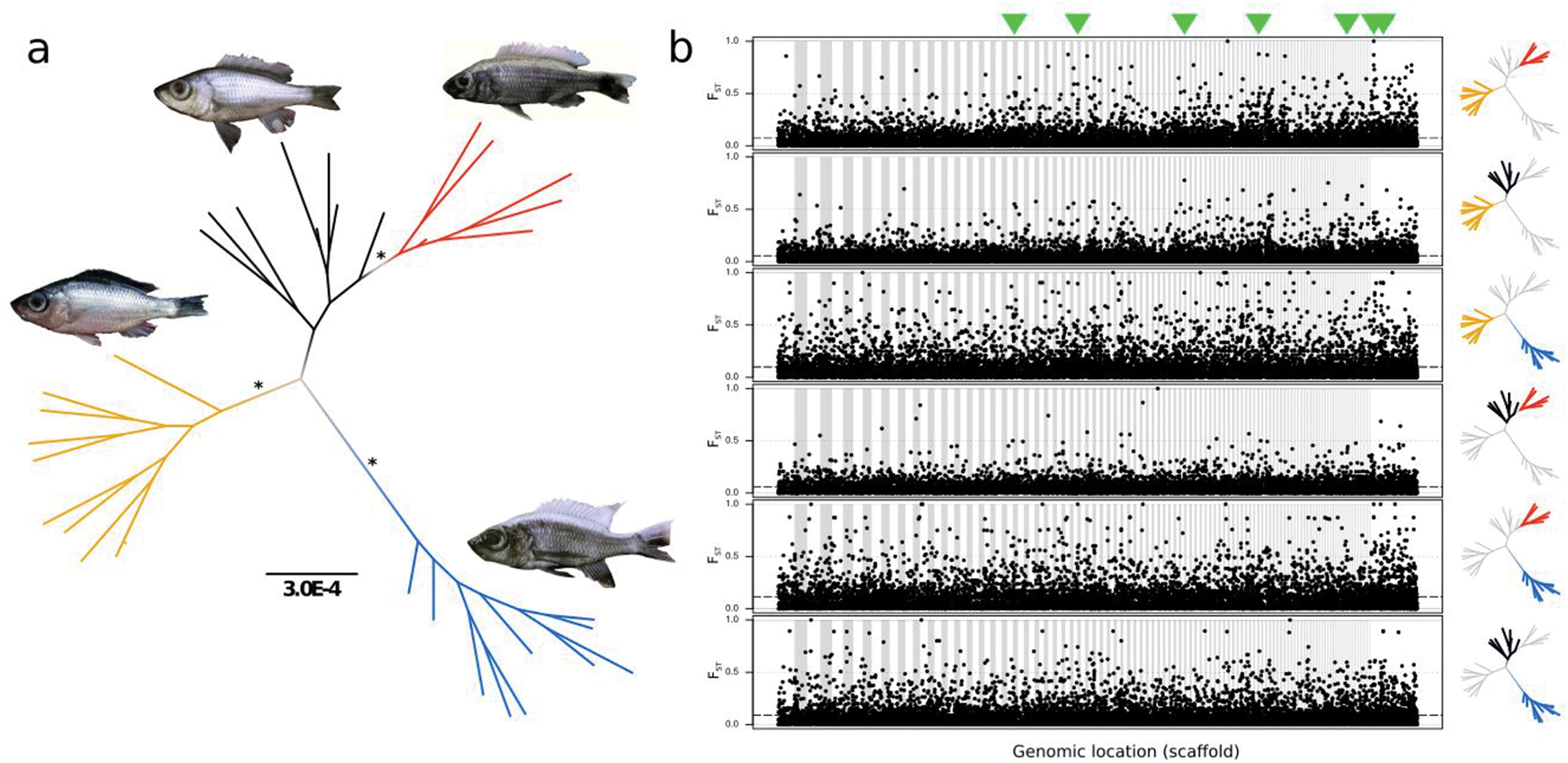
(a) Maximum Likelihood tree (unrooted) of phylogenomic structure among *Diplotaxodon* species. Yellow - *D*. ‘macrops black dorsal’; red - *D*. ‘limnothrissa black pelvic’; black - *D*. ‘macrops offshore’; blue - *D*. ‘macrops ngulube’. Asterisks indicate > 95% bootstrap branch support. Scale bar indicates genetic divergence (nucleotide divergence per site). (b) Genome wide pattern of pairwise F*_ST_* divergence between populations. Scaffolds with minimum length of 100kb containing a minimum of 10 SNPs are displayed. Individual scaffold boundaries are indicated by alternate white/grey background. Dashed lines indicate the global pairwise F*_ST_* average. Highlighted groups in the phylogenetic trees on the right hand side indicate the population pairs. Green arrow heads on top of the figure indicate locations of candidate regions supported by all three candidate outlier approaches (see Figure S3).

### Candidate genomic regions under selection

We applied three independent outlier detection approaches, which highlighted 242 loci (2.1% of total), distributed across 103 genomic scaffolds of the *Maylandia* (*Metriaclima*) *zebra* reference genome v1.1 ^32^. The number of loci highlighted by individual methods ranged from 125 (1.1 % of total, *Stacks*) to 96 (0.8 % of total, *Bayenv*). Of the total candidate loci, 189 (77.8 %), 41 (16.9 %) and 13 (5.3 %), were highlighted independently by one, two, or all three methods, (see Figure S3) and these were assigned to 134, 26 and 8 candidate regions on 103, 26 and 8 genomic scaffolds, respectively. Table 1 summarizes the location and characterizes the gene complements of the characterizes the gene complements of the 26 genomic regions supported by at least two26 genomic regions supported by at least two independent outlier identification approaches (Table S3 lists gene models associated with the candidate regions on the 103 genomic scaffolds).

**Table 1:**
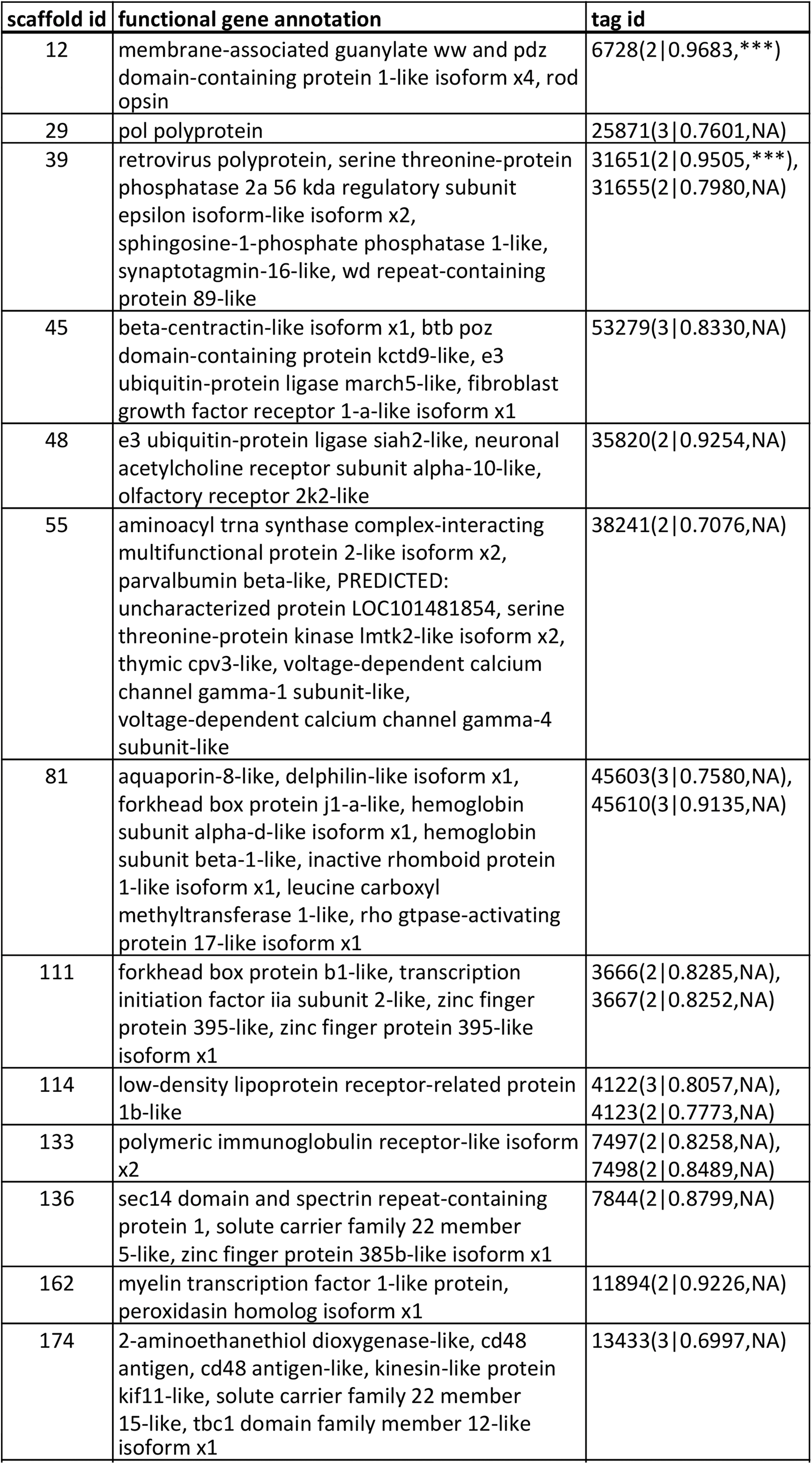
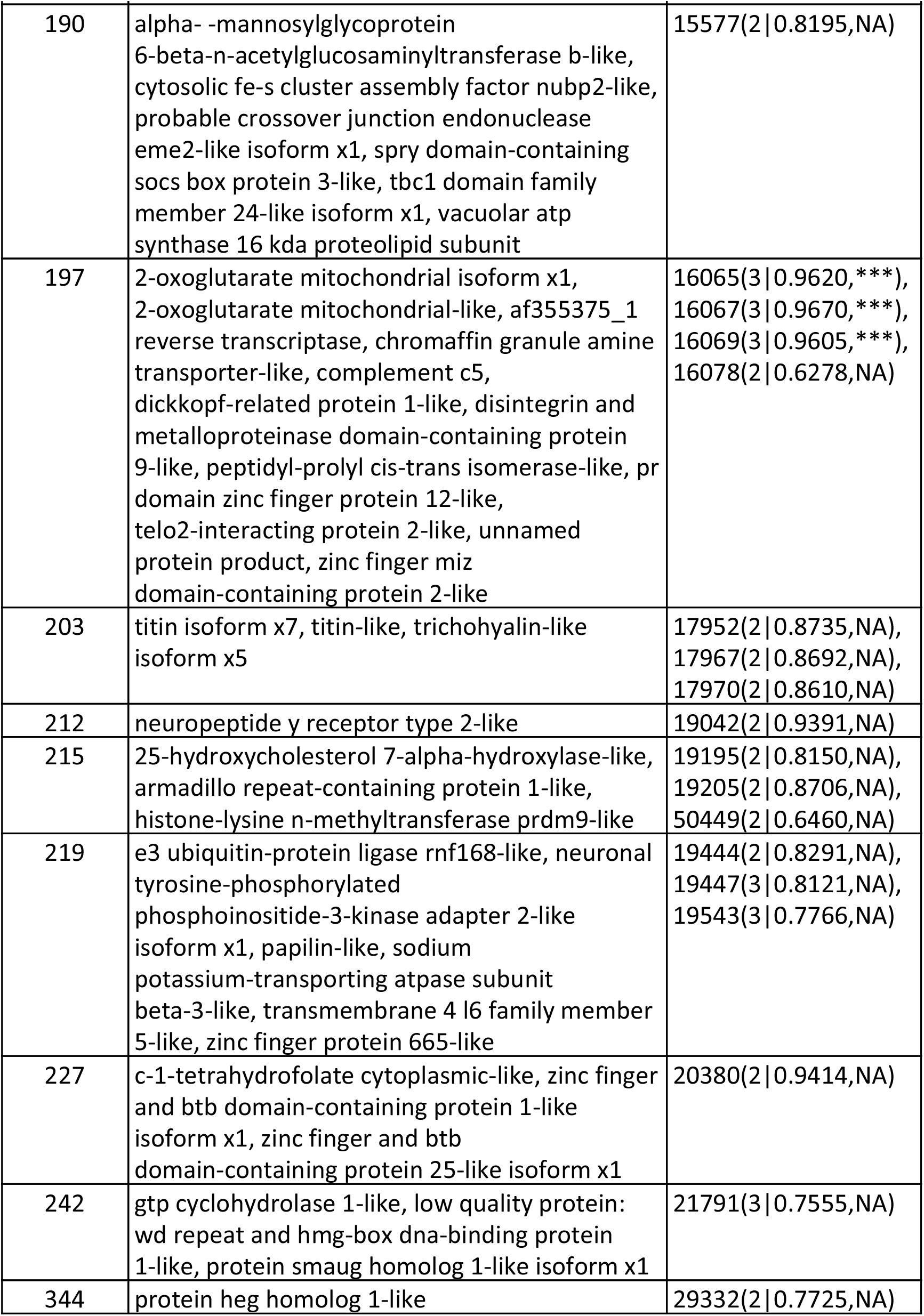
Genomic location (scaffold id), putative functional annotation of genes (if available) and RADtag ids localized in candidate regions under selection. Numbers in parenthesis after the RADtag id describe the number of independent outlier detection approaches supporting the respective tag, followed by the same tag’s ARR and smoothed ARR level of significance (NS: > 0.05; *: 0.05-0.005; **: 0.005-0.001; ***: <0.001), inferred by the eye morphology informed analysis. Displayed are only gene complements for regions supported by at least two independent outlier detection approaches. Full list of highlighted regions including gene ids can be found in supplementary table S3.

Gene ontology enrichment analyses of genes in the candidate regions under divergent selection highlighted GO terms associated with sensory perception (e.g. axon extension, neuron development) and photoreceptor development (dynactin complex), embryonic development and morphogenesis (fibroblast growth factor receptor signalling pathway, positive regulation of cell proliferation, developmental growth involved in morphogenesis), and oxygen binding/transport (haemoglobin complex, oxygen transport/binding, gas transport, haem binding). Specific candidate genes under divergent selection associated with visual perception, and signal transduction include *ACTR1B* (Beta centractin), *Rh1* (Rhodopsin), *PPIase* (Peptidyl-prolyl cis-trans isomerase), *SGPP1* (Sphingosine-phosphate 1 phosphatase 1), and *DENND4B* (see Tables 1 and S5 for details). The phosphatase coded by *SGPP1* catalyses the degradation of sphingosine-1-phosphate, a key regulator in photoreceptor development ^33,34^. With respect to putative physiological adaptations to deep-water environments, a genomic region centred around two haemoglobins, HBA (haemoglobin subunit alpha) and HBB1 (haemoglobin subunit beta-1), was highlighted by all three candidate outlier approaches.

Further candidates under putative selection include genes central for craniofacial and eye morphogenesis, such as *ALX3* (Aristaless-like 3), *PXDN* (Peroxidasin), *FOX* (Forkhead box transcription factor, 2x), *ZMIZ* (Zinc finger miz domain containing protein, 2x), *RPGRIP1L* (retinitis pigmentosa GTPase regulator interacting protein 1, synonym Fantom), *MEIS2* (homeobox meis2 protein), *FGFR1* (fibroblast growth factor receptor 1), *SOX* (Sry box transcription factor, 2x), *DKK1* (Dickkopf1), *NEUCRIN* (Draxin) and *TCF7L1* (transcription factor 7-like 1, formerly known as *TCF3*). *RPGRIP1L* has been shown to interact biochemically with *RPGR* (Retinitis Pigmentosa GTPase Regulator) ^35^, which plays a central role in controlling access of both membrane and soluble proteins to the photoreceptor outer segment. Loss and mutations in *RPGR* have been associated with a range of retinal diseases in human patients, including a variant of cone dystrophy, characterized by progressive dysfunction of photopic (cone-based) day vision with preservation of scotopic (rod-based) night vision ^36^. *TCF7L1* is involved in the regulation of early embryonic craniofacial development via the Wnt/β-catenin signalling pathway and is expressed during human embryonic eye development ^37^. Generally the Tcf/Lef family of molecules mediate canonical Wnt signaling by regulating downstream target gene expression ^38^. *TCF7L1* has been demonstrated to directly repress *SOX4* ^39^ which is expressed in early zebrafish eye development ^40^ and also plays an active role in Wnt signalling by stabilizing β-catenin via a complex feedback loop ^41^. Knockdown of *SOX4* in zebrafish resulted in structural malformations of the eye ^40^. *SOX2*, also affects Wnt signalling via feedback inhibition, and has been shown to crucially regulate retina formation in *Xenopus* ^42^ and mice ^43^. Mutations in the underlying gene have been associated with recessively inherited frontonasal malformation in humans ^44^. Within the candidate regions identified by our analyses we found cases of co-localization of genes potentially functionally relevant for deep-water adaptation, such as the close proximity of *ZMIZ*, *DKK1* and *PPIase* (scaffold 197, Figures 3c and S3), and *ACTR1B*, *FGFR1* and *TCF7L1* (scaffold 45, see figure S3). *ACTR1B* is involved in regulating photoreceptor cell differentiation ^45^ and survival ^46^*. ZMIZ* has been previously shown to directly interact *in vitro* with *Msx2*, an important regulatory element involved in skull- ^47^ and specifically in eye morphogenesis ^48^. *DKK1* is an antagonistic inhibitor of the Wnt/β-catenin signalling pathway, repeatedly implicated as a central mediator of craniofacial development in vertebrates ^49,50^, including cichlids ^51,52^. In Lake Malawi cichlids an amino acid substitution in β-catenin has been found to be associated with alternate jaw morphologies. *DKK1* inhibits the stabilization of β-catenin by binding to *LRP6*. Colocalized with these two factors is a peptidyl-prolyl cis-trans isomerase (*PPIase*). Generally, *PPIases* are ubiquitous proteins, but in the context of this study it is worth noting that members of a *PPIase* subgroup (cyclophilins) have been shown to play a critical role in opsin biogenesis in *Drosophila* ^53^ and cattle ^54^. For some of the above mentioned candidate genes our outlier approaches have highlighted more than one paralog on separate genomic scaffolds. *ZMIZ*, *FOX* and *SOX* transcription factors were present in two separate candidate regions, each (see tables 1 and S3).

### Genomic regions associated with interspecific eye size variation

Both vertical (Figure 3a) and horizontal (Figure S5) eye diameter differed significantly among the four *Diplotaxodon* species (ANOVA: vertical eye diameter F_3,171_ = 104.6, P < 0.001; horizontal eye diameter F_3,171_ = 127.7, P < 0.001;), with *D. limnothrissa* eye diameters being consistently significantly smaller in all pairwise comparisons (TukeyHSD: P < 0.001, Table S4). Enlargement of the eyes is often associated with adaptation to deep-water environments ^13,14^. We applied a Bayesian linear model approach using the observed eye size differences to identify genomic regions particularly associated with this phenotypic trait. These analyses highlighted the population allele frequencies of 42 loci (0.4 % of total, Figure 3b), clustering into 37 genomic windows, as being highly correlated with interspecific eye diameter differences, indicated by their average relative rank (ARR >= 0.95) across 20 independent Bayenv runs. Eight SNPs (0.07 % of total) were found to be highly significantly associated with eye diameter using our most stringent filtering criteria (ARR >= 0.95 and smoothed ARR p < 0.001). This set of candidate SNPs were in six genomic regions (Figure 3c). Three of these six genomic regions inferred as most significantly correlated with interspecific eye diameter variation, are also highlighted as candidate regions under selection by at least two of three independent outlier detection approaches (Figure 3c). GO enrichment analyses of the gene complements identified by the eye diameter-informed analyses indicates significant overrepresentation of GO terms associated with e.g. ‘regulation of response to external stimulus’ and ‘intermediate filament organization’. The corresponding genomic regions contain genes encoding for the photopigments, *Rh1*, *OPN4* (melanopsin) and *OPN5* (neuropsin), as well *SGPP1 and PPIase*, a structural eye lens protein (*BFSP2*), genes expressed during normal eye development (*TMX3, XFIN*), and central transcription factors regulating embryonic craniofacial development (*ZMIZ, DKK1, ALX3*). Particularly *BFSP2*, *OPN4*, *TMX3* and *XFIN* were found in close proximity (scaffold 215). Tables 2 and S5 detail the gene complements in the genomic regions most significantly associated with eye size differences. Candidate genes contained in regions highlighted by both the eye morphology informed analysis and the candidate outlier detection approaches include *Rh1*, *SGPP1*, *ZMIZ*, *DKK1*, *PPIase*, *ALX3* and *RPGRIP1L* (Table S5).

**Table 2:** Genomic location (scaffold id), putative functional annotation of genes (if available) and RADtag ids localized in candidate regions most significantly associated with interspecific eye size variation. Numbers in parenthesis after the RADtag id describe the tag’s ARR and smoothed ARR level of significance (NS: >0.05; *: 0.05-0.005; **: 0.005-0.001; ***: <0.001). Displayed are only gene complements for regions supported at significance level p<0.001. Full list of highlighted regions supported by ARR >= 0.95 including gene ids can be found in supplementary table S5.

| scaffold id | functional gene annotation | tag id |
| --- | --- | --- |
| 12 | membrane-associated guanylate ww and pdz domain-containing protein 1-like isoform x4, rod opsin | 6728(0.9683,***) |
| 39 | a-kinase anchor protein 6, neuronal pas domain-containing protein 3-like, serine threonine-protein phosphatase 2a 56 kda regulatory subunit epsilon isoform-like isoform x2, sphingosine-1-phosphate phosphatase 1-like, synaptotagmin-16-like, wd repeat-containing protein 89-like | 31602(0.9562,*),<br>31651(0.9505,***) |
| 159 | dynein heavy chain axonemal-like | 11119(0.9666,***) |
| 197 | 2-oxoglutarate mitochondrial isoform x1, 2-oxoglutarate mitochondrial-like, af355375_1 reverse transcriptase, chromaffin granule amine transporter-like, dickkopf-related protein 1-like, disintegrin and metalloproteinase domain-containing protein 9-like, peptidyl-prolyl cis-trans isomerase-like, zinc finger miz domain-containing protein 2-like | 16065(0.9620,***),<br>16067(0.9670,***),<br>16069(0.9605,***) |
| 215 | afg3-like protein 2-like, c-c chemokine receptor type 9-like, chromodomain y-like isoform x2, fas-activated serine threonine kinase, kelch-like protein 18-like isoform x1, melanopsin-like, nucleolar transcription factor 1-like, phakinin-like, poly -binding-splicing factor puf60-like isoform x1, protein disulfide-isomerase tmx3-like, zinc finger protein xfin-like isoform x1 | 19268(0.9664,***) |

**Figure 3.**
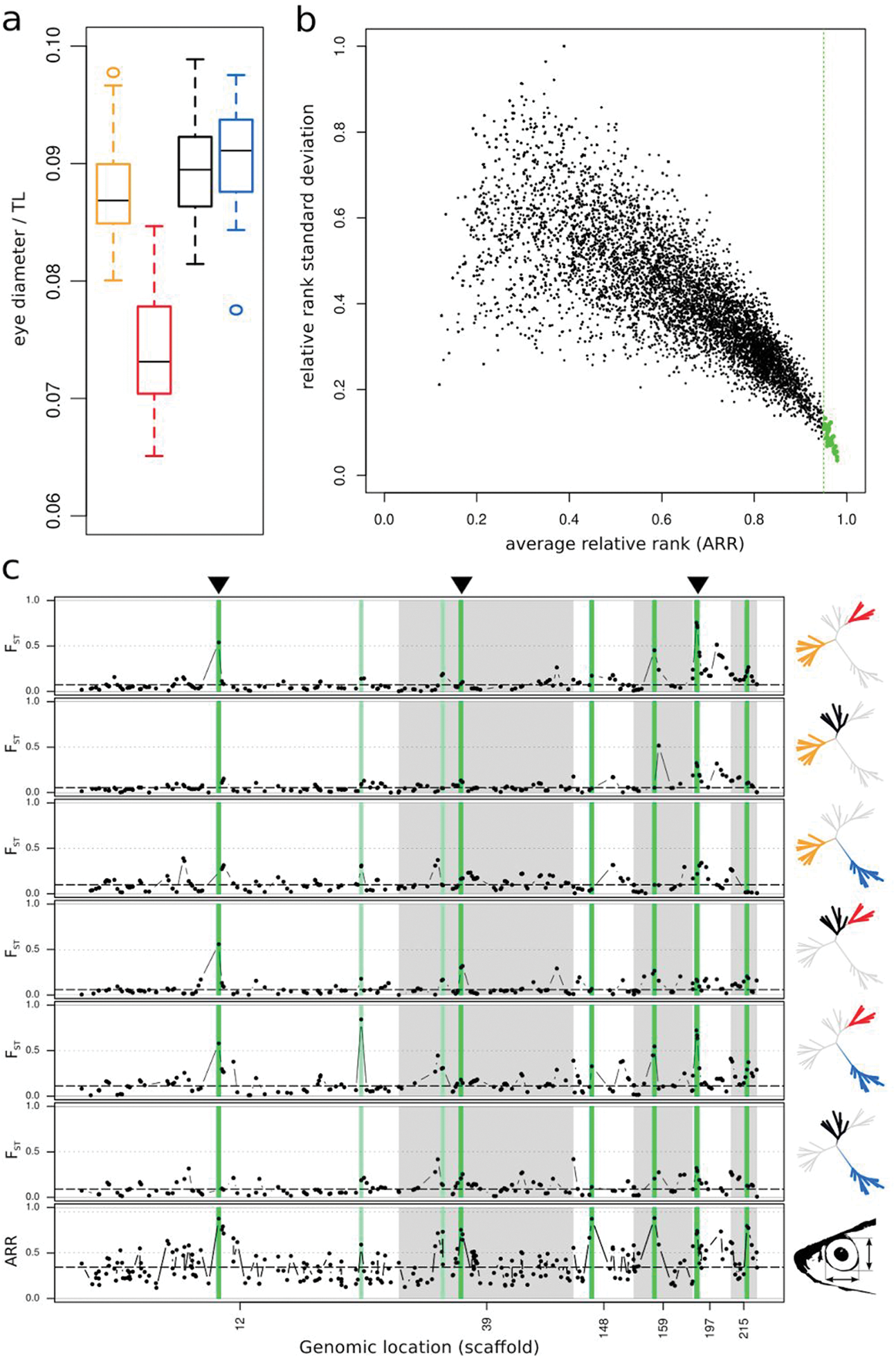
(a) *Diplotaxodon* eye diameter (normalized by total length of the fish, TL) across the four species. Yellow - *D*. ‘macrops black dorsal’; red - *D*. ‘limnothrissa black pelvic’; black - *D*. ‘macrops offshore’; blue - *D*. ‘macrops ngulube’. (b) Pope plot, summarizing correlations of population allele frequency with vertical eye diameter, inferred from 20 independent Bayenv runs. Dots illustrate per locus average relative rank (ARR) versus relative rank standard deviation. The green vertical line delimits the 95th ARR percentile. The 42 loci most consistently correlated with eye size (ARR >= 0.95) are shown in green. (c) Pairwise F*_ST_* divergence (top six panels) and global allele frequency correlations with interspecific vertical eye diameter differences (bottom panel). Displayed are the six scaffolds containing the most significantly correlated loci. Corresponding regions are highlighted in shades of green (lightgreen - ARR >= 0.95; darkgreen - ARR >= 0.95 and smoothed ARR *P* < 0.001). Dots represent the kernel smoothed averages across 50kb windows. Dashed lines indicate the genome wide average of F*_ST_*/ARR. Population pairs are indicated by the highlighted populations in the phylogenetic trees on the right hand side. Black arrowheads on top indicate regions that were also supported as candidate outlier loci by at least two of three independent outlier detection approaches. Tables 2 and S5 summarizes the gene complements in highlighted regions.

### Functional adaptations in candidate genes

Using genome-level resequencing data obtained via the Malawi cichlid diversity sequencing project (NCBI Bioproject PRJEB1254), we examined nucleotide polymorphisms to identify potentially functional substitutions in a number of genes highlighted by the population level analyses described above, specifically the genes coding for rhodopsin, phakinin, melanopsin, as well as haemoglobin subunits alpha and beta. The resequencing data include 156 individuals from 78 haplochromine species from in and around Lake Malawi, including six *Diplotaxodon* species. These analyses revealed a number of non-synonymous substitutions, as well as indel-, 5’- and 3’-UTR polymorphisms, potentially relevant for visual and physiological adaptation to deep-water conditions (Table S6). Across *Diplotaxodon* species, we identified a total of eight non-synonymous and three 3’-UTR polymorphisms in the RH1 gene coding for rhodopsin. Three of the non-synonymous polymorphisms involve variants so far restricted (private) to *Diplotaxodon*, i.e. were not observed in any of the other samples from the greater Lake Malawi species flock. One further non-synonymous variant appears private to the *Diplotaxodon-Pallidochromis* lineage. Three of the amino acid residues affected by non-synonymous substitutions (amino acid positions 83, 133 and 189) were previously considered likely to be involved in spectral tuning (see discussion for further details). Variants private to *Diplotaxodon*, were also detected in the genes coding for phakinin and melanopsin (Table S6). We detected a total of 13 amino acids containing non-synonymous polymorphisms in the haemoglobin subunit beta gene. One of the variants appeared private to *Diplotaxodon*, while several others were observed in high frequency in *Diplotaxodon* and in additional taxa of a distinct deep-water lineage of benthic Lake Malawi cichlids, including *Alticorpus*, *Lethrinops* and *Aulonocara*. Several of the affected residues are associated with changes in O_2_ affinity in the human haemoglobin subunit beta homologue.

## Discussion

We identified genomic regions showing signals of strong differentiation between four species of deep water *Dipotaxodon* species in Lake Malawi. In shallow water cichlids, species tend to segregate in habitat, diet and depth distributions. Thus, we predicted such resource partitioning is likely to be taking part in deep water cichlids, and that genomic regions associated with adaptation should appear overrepresented in outlier loci. Gene ontology enrichment analyses highlighted GO terms associated with sensory perception and photoreceptor development, embryonic development and morphogenesis, and oxygen binding/transport. We first used RADseq data, and assumed regions within 50kb of the highlighted SNP loci were candidates for selection among *Diplotaxodon* species. We subsequently used genome-level resequencing data and found that patterns of SNP diversity in coding (and regulatory) regions may reflect ecological differences between *Diplotaxodon* and the rest of the species flock.

Our analyses indicate a role for key genomic regions containing genes which could be associated with adaptation to depth. A region centred around two haemoglobin genes shows a signal consistent with selection, and this region is relevant for physiological adaptation to deep-water conditions in two ways. The first is through enhancing the “Root effect”, a pH dependent decrease in oxygen-carrying capacity of some fish haemoglobins. This facilitates O_2_ secretion into the swim bladder, the specialized organ found in most teleost fishes used to achieve neutral buoyancy in open water by regulating partial gas pressure in response to the ambient hydrostatic pressure. At the molecular level, such ‘Root haemoglobins’ are characterized by a range of amino acid replacements in the globin α- and β-chains, compared to ‘normal’ haemoglobins ^55–57^ and may have evolved independently a number of times ^58^. The two affected *Diplotaxodon* haemoglobin subunit genes fulfil the minimal structural requirements for Root haemoglobins as previously defined ^59^. The second way selection could act on this region is via haemoglobin O_2_ binding affinity Given teleost fishes can differ in their tolerance towards low levels of dissolved oxygen, and the amount of dissolved oxygen in water typically decreases with depth, it is plausible that divergent selection is operating on oxygen tolerance in Lake Malawi ^61^. Meromictic freshwater lakes exhibit complete oxygen depletion below a defined oxic-anoxic boundary layer, which in Lake Malawi has been identified at ~230 m water depth ^62^. Selection for high O_2_ affinity haemoglobin alleles was previously demonstrated in response to altitude related hypoxia in birds ^63–65^. A number of the residues affected by non-synonymous variants in *Diplotaxodon* (Table S6) have been associated with changes in oxygen affinity in the human haemoglobin subunit beta-1. While both the Root effect and haemoglobin oxygen binding affinity have previously been predicted to be likely targets of natural selection ^2,61^, the current study is, to our knowledge, the first to find evidence for selection associated with haemoglobin genes in fish. The exact effect of the observed non-synonymous changes on the Root effect and/or the O_2_ affinity of *Diplotaxodon* haemoglobins needs to be determined experimentally, but the presence of the same changes in other unrelated deep-water Lake Malawi genera such as deep-benthic *Lethrinops* and *Alticorpus* spp. is consistent with adaptation to depth. Whether these changes have arisen multiple times independently, or been acquired through introgression warrants further investigation. There is evidence of hybridization as a driver in adaptive radiation in the Lake Malawi species flock ^66–68^ and these loci could allow a test of whether these deep water adaptations have been retained after a hybridization event, and allowed subsequent radiation into this challenging habitat ^67^

Previous work has shown that the four *Diplotaxodon* species we studied differ significantly with respect to head- and overall body morphology ^27^, and craniofacial variation in cichlids is frequently associated with trophic adaptations. Strongly differentiated cranial dimensions include eye size differences and light conditions at different water depths are a likely selective driving force. We identified candidate genomic regions under selection that are highly enriched for genes involved in the regulation of craniofacial development. Our F*_ST_* outlier detection approaches highlight a number of *Wnt* factors, further supporting the central role of *Wnt* signalling for regulating cichlid craniofacial gene expression ^69^. One genomic region in particular (scaffold 197, Figure 3b) received strong support in all analyses, and contains a group of colocalized genes; namely *ZMIZ* (skull and eye morphogenesis), DKK1 (vertebrate craniofacial development) and a PPIase (opsin biogenesis). The close proximity of *ZMIZ* and *DKK1* in particular may indicate they are inherited together as a craniofacial “supergene”.

The larger eye of species in the *Diplotaxodon macrops* “complex” relative to those in the *Diplotaxodon limnothrissa* “complex” is consistent with their presumed depth distributions (Figures <u>3a</u> and S5). The environment at depths of 50 - 200 m ^70^ is depauperate of long wavelength light and dominated by shorter wavelength blue light as depth increases ^12^. Our analyses consistently highlight a genomic region centred around the *Rh1* gene (scaffold 12, Figure 3b), which codes for rhodopsin, the principal photopigment of retinal rod photoreceptors, central for scotopic vision under dim light conditions. Changes in both the coding sequence as well as in gene expression of opsins ^71^ may mediate visual performance across light environments, and the pivotal role of rhodopsin for visual adaptation in deep water light environments has been confirmed by a number of studies ^6,19,25,72,73^. Whether the observed non-synonymous variants in the rhodopsin gene (Table S6) result in functionally important shifts of the spectral sensitivity of the photopigment between *Diplotaxodon* species needs further investigation. However, it is worth noting that three of the observed 10 amino acid residues found affected by non-synonymous polymorphisms identified in the current study (amino acid positions 83, 133 and 189) have previously been considered likely to be involved in spectral tuning, because of their close proximity to the chromophore or the chromophore-binding pocket ^25^. Amino acid replacements at position 83, specifically, have been demonstrated experimentally to cause spectral shifts towards blue ^25^. A further four affected residues (166, 169, 297, 298) have also been implicated in a recent study of spectral shifts via mutations in the rhodopsin gene in an isolated East African crater lake ^21^.

Our eye-size informed analyses suggest that *BFSP2*, the gene coding for phakinin, has been diverging among *Diplotaxodon* species. The lens of the vertebrate eye is composed of specialized epithelial lens fibre cells containing beaded filaments, specific cytoskeletal structures unique to the lens ^74^. Phakinin is one of the two principal proteins forming the beaded filaments, so our results also suggest a role for eye lens structure in adaptation to dim, short wave-length light environments, a concept that has so far attracted very little attention. The role of beaded filaments in lens biology is not fully understood, but they appear essential in maintaining optical clarity and transparency of the lens ^75,76^. Mutations in *BFSP2* are associated with cataract formation, i.e. a clouding of the eye lens in humans ^77,78^. Cataracts reduce the intensity and alter the chromaticity of light traveling through the lens ^79^, with potentially great effect on visual perception ^80^. After cataract surgery patients usually report a change in color appearance associated with additional short-wavelength light reaching the retina ^79,80^. The wider implication is that *BFSP2* may specifically regulate lens transparency for blue light and could be a key- and previously unrecognised mediator for adaptation in dim, blue-dominated light environments.

*BFSP2* is co-localized with *OPN4*, *TMX3* and *XFIN*. The role of the latter genes in visual adaptation is unknown. However, *XFIN* is expressed during the formation of the retina in *Xenopus* ^81^ and deletion of the *TMX3* gene in humans has been linked to a genetic disease associated with retarded growth of the eye ^82^. *OPN4* codes for the opsin based photopigment melanopsin, which is central to a distinct photoreceptor class, the melanopsin retinal ganglion cells (mRGCs), that was discovered only relatively recently ^83^. Initially mRGCs were shown to mediate so-called non-image forming visual responses ^84^, but more recent evidence suggests that mRGCs may contribute significantly to assessing brightness and play a more general role in supporting vision in mammals ^85^ and it may well play a role in deep-water cichlid vision. In Lake Malawi cichlids the associated genomic region is clearly highly enriched for vision related genes and might represent another coadapted gene complex. Exploration of whole genome resequencing data obtained for the larger Malawi flock revealed non-synonymous mutations private to the deep water lineage in both the *BFSP2* and the *OPN4* gene, as well as intron and 5’-UTR variation in the respective genes between *Diplotaxodon* species. Further investigation is required to fully understand how this genomic region may be involved in deep-water visual adaptation. The potential role of lens structure in adaptation is further confirmed by the highlighting of the *FGFR1*, which is considered essential for lens fibre differentiation ^86^ and *MEIS2*, a gene that directly regulates *Pax6* during vertebrate lens morphogenesis. The latter transcription factor has been demonstrated to play essential roles in lens differentiation and has previously been referred to as the ‘master control gene for morphogenesis and evolution of the eye’ ^87,88^.

Depth- and habitat segregation may be caused by a number of factors including competition for resources, breeding territories or enemy-free space ^89^. The four *Diplotaxodon* also differ in male nuptial coloration ^27^ which strongly implies an important role of visually informed mate choice, despite the twilight conditions they experience in their natural environments. While many pelagic fish use counter-shading in background matching for camouflage ^90^, *D. ‘*macrops ngulube’ and *D. ‘*macrops black dorsal’ males have an opposite nuptial coloration which may allow them to be more visible. The sensory drive hypothesis^91^ predicts that visual systems (along with signals and signalling behaviour) will differentiate if local environments differ in their signal transmission quality. The observed interspecific differences in male nuptial colour in *Diplotaxodon* in combination with the inferred genomic footprints of sensory adaptation are consistent with the idea that reproductive isolation could arise as a consequence of sensory drive in deep water systems.

In summary, our work has shown that the selection pressures associated with deep water environments can be identified by their effect on the genome. In addition to genes previously associated with depth related spectral shifts (rhodopsin), we identify novel mechanisms of adaptation to deep water conditions worth further investigation (e.g. Root effect haemoglobins and eye lens filament proteins). Outlier tests tend to highlight large-effect loci with relatively simply genetic architecture ^92^ so these will be interesting possibilities with which to identify parallel evolution in other systems such as populations experiencing high altitude hypoxia, or marine deep water systems. Our results provide evidence of fixed genomic changes since deep water conditions in Lake Malawi were attained possibly as recently as 75,000 years ago ^28,29^, and raise the intriguing possibility that hybridization between *Diplotaxodon* and the deep-benthic clade of cichlids may have facilitated the latter’s expansion into the twilight zone. Finally, we find that in candidate genomic regions under selection functionally associated genes are frequently in close proximity, such that cichlid adaptations and ecological differentiation may be facilitated by the presence of linked, coadapted gene complexes, or “supergenes”.

## Methods

### RAD data sampling, DNA extraction, library preparation and Illumina sequencing

A total of 40 individuals from four *Diplotaxodon* species were collected from Nkhata Bay (Figure 1, Table S1), photographed, and fin clips stored in ethanol at −80℃. Genomic DNA was extracted from fin clips using the Qiagen DNAeasy Blood and Tissue Kit, according to manufacturer’s instructions. DNA was quantified using PicoGreen fluorimetry (Quant-iT PicoGreen Kit, Invitrogen) and quality checked on 0.8% agarose gels. Paired-end RAD libraries were prepared by the NERC/NBAF facility at Edinburgh Genomics, following ^93^ and ^94^Genomic DNA was digested using *SbfI*, a barcoded RAD P1 adapter ligated, followed by sonic shearing, size selection and ligation of P2 adapters. Libraries were PCR amplified, quantified and sequenced in separate flow cells on an Illumina HiSeq 2000 platform with 100 bp, paired-end chemistry.

### RAD data processing

Modules from the Stacks v.1.20 ^95^ program suite were used for the initial processing of the RAD data as follows: Raw reads were demultiplexed based on their in-line barcode and quality trimmed using the program *process_radtags*. Putative PCR-duplicates were subsequently removed using the program *clone_filter* from Stacks v.1.20. The remaining reads for each individual were mapped to the draft genome of *M. zebra* ^32^, version MetZeb1.1_prescreen (downloaded from the link provided; last accessed 23.12.2016) http://archive.broadinstitute.org/ftp/pub/assemblies/fish/M_zebra/MetZeb1.1_prescreen/M_zebra_v0.assembly.fasta; using BWA v.0.7.10-r789 ^96^, allowing for up to 8 mismatches per read. Any reads mapping to more than one genomic location were removed from the dataset using a custom script (*split_sam.pl*) and the remaining reads were converted to bam format using Samtools v.0.1.19-4428cd ^97^. The program *pstacks* from Stacks v.1.20 was used to group the mapped reads into stacks (minimum of 5 identical reads per stack) and call SNPs. A global catalog of RAD tags was then built using *cstacks* from all individuals with at least 20,000 valid tags identified by *pstacks* (individuals with fewer valid tags were omitted from further analyses). The *populations* program from Stacks v.1.20 was used to calculate basic population genetics statistics and to produce data files for downstream analyses (e.g. in plink format, structure format) for each population individually, for each pairwise population comparison and across a global dataset of all populations. Analyses were limited to tags identified in all populations and in at least 80% of individuals per population, respectively. Maximum likelihood inference was performed on the concatenated RAD tags using RAxML v8.2.4 ^98^. SNP datasets to be used in downstream analyses were limited to only a single SNP per RAD-tag (*--write_single_snp* option in *populations*) to reduce the effects of linkage. We furthermore excluded any singleton SNPs, i.e. sites with minor allele counts of one, from any global analyses. Further conversions to file formats not supported by *populations* were performed by PGDSpider v.2.0.7.3 ^99^ or by custom scripts. Principal component analysis and discriminant analysis of principal components ^100^ was performed using the Adegenet package in R ^101^.

### Identification of putative candidate loci under selection

Three independent approaches to outlier detection were applied: (1) Pairwise Fst values were calculated by *populations*. A Fisher’s exact test as applied by the sliding window algorithm implemented in *populations* (window size 50kb) was used to calculate significance levels ^95^ and loci with p < 5e-7 in at least one pairwise comparison were considered candidate loci under selection identified by *Stacks*. (2) The Bayesian approach to outlier loci detection implemented in *Bayescan* v.2.1 ^102^ was applied to pairwise, as well as a global SNP dataset, produced by *populations*. In all Bayescan analyses, prior odds for the neutral model were set to 10, with the remaining parameters set to default. A false discovery rate (FDR) threshold of 0.05 was applied and loci with α < 0.05 in at least one pairwise comparison or the global Bayescan run were considered candidate loci identified by *Bayescan*. (3) *Bayenv* v.2 ^103^ was used to identify candidate outlier based on the global SNP dataset obtained for all four populations. *Bayenv* applies a Bayesian linear model method that accounts for population history by incorporating a covariance matrix of population allele frequencies and accounting for differences in sample size among populations ^103^. 10 independent covariance matrices were constructed for sets of 5,000 SNPs randomly selected from the global dataset as represented in a vcf file produced by *populations*, using a custom script (*vcf_2_div.py*). In brief, for each random set covariance matrices were obtained by running *Bayenv* for 100,000 iterations. Convergence of covariance matrices was assessed visually in R ^104^ and a final covariance matrix was obtained by averaging across the 10 independent runs. Using this matrix to account for population history *Bayenv* was then run for 20 independent replicates with 1,000,000 iterations each on a set of SNPs present in at least 5 individuals in each population. Candidate loci under selection were identified based on the population differentiation statistic *xTx* generated by *Bayenv* as follows: For each run we ranked loci based on their respective value for *xTx*, and calculated a rank statistic similar to the empirical p-value approach used by ^105^. In our approach the highest ranking SNP would be assigned the highest relative rank of 1, while the lowest ranking SNP would be assigned a rank of *1/N*, with *N* being the total number of SNPs in the dataset. For each SNP we then calculate the average relative rank (ARR) and standard deviation across the 20 independent *Bayenv* runs. Loci yielding an ARR in the 95^th^ percentile were considered candidate loci under selection identified by *Bayenv*. Genomic candidate regions were defined as +−50kb windows up and downstream of candidate SNPs, supported by one, two, or three outlier detection approaches, respectively. Candidate regions of consecutive candidate SNPs were merged in case an overlap between the corresponding windows was detected.

### Identification of SNPs correlated with morphological differences between populations

The Bayesian linear model approach implemented in *Bayenv ^103^* is frequently used to infer correlations between SNP allele frequencies and environmental variables. The method yields Bayes factors (BF), which are interpreted as the weight of evidence for a model in which an environmental factor is affecting the distribution of variants relative to a model in which environmental factors have no effect on the distribution of the variant ^105^. The four *Diplotaxodon* populations analysed in the current study have been previously identified to differ significantly in head morphological traits ^27^, which in turn are known to correlate with environmental variables in cichlids ^106^. In particular, *Diplotaxodon* ‘limnothrissa black pelvic’ has a smaller eye than the other species. We applied the *Bayenv* approach, using vertical and horizontal eye diameter (normalized by total fish length) from ^27^, to identify SNPs correlated with eye morphological differences between populations and to characterize the genomic regions involved in regulating the observed morphological differences. The full *Bayenv* workflow and the subsequent analyses are made available as Jupyter notebooks in a dedicated Github repository. Prior to *Bayenv* analyses population averages were standardized by subtracting the global mean and dividing the result by the global standard deviation. Standardized population averages were then used as ‘environmental variables’ in 20 independent runs of *Bayenv*, each using 1,000,000 iterations. Transformed rank statistics by means of average relative ranks (ARR) of Bayes factors were calculated across the 20 runs as described above. We then applied a Gaussian weighting function to generate a kernel-smoothed moving average of the transformed rank statistic for each polymorphic site (based on 100 kb sliding windows centered at each SNP). To test for statistical significance of windows we applied a bootstrap resampling procedure (10,000 permutations). In each permutation new values for ARR were sampled with replacement from across the dataset and the smoothed statistic was calculated for each replicate set using the coordinates of the original SNP for the weighing function. For each SNP, the data obtained via bootstrap resampling was used as empirical null distribution of the test statistic against which the original smoothed average ARR was compared to determine a P-value. The approach we have implemented is similar to the method used by the *populations* program of the *stacks* software suite ^95^. Windows significant at the p<0.001 level were considered candidate regions associated with eye morphological differences if they also contained at least one individual SNP locus yielding an ARR in the 95^th^ percentile of the original distribution.

### Identification the function of genomic regions under selection

Gene models for *M. zebra* ^32^ were obtained from the Broad Institute (downloaded from ftp://ftp.broadinstitute.org/pub/vgb/cichlids/Annotation/Protein_coding/; last accessed 23.12.2016). Peptide sequences were subjected to a similarity search against a custom build of the NCBI’s nr protein database (restricted to Metazoan proteins) using BLAST ^107^ and screened for known domains using InterProScan 5.8-49.0 ^108^. Results from these analyses were reconciled in Blast2GO v.3 ^109^ to obtain putative functional annotation including Gene Ontology (GO) terms ^110^ for *M. zebra* gene models, where possible. GO term enrichment analyses using Fisher’s exact tests with multiple testing correction of FDR ^111^, as implemented in Blast2GO v.3 ^109^, were performed for gene complements in genomic candidate regions supported by one, two, or three candidate outlier approaches, respectively.

### Whole genome resequencing data

Following up on our initial results, we selected the genes coding for rhodopsin, phakinin and melanopsin, as candidate genes for visual adaptations, as well as the genes coding for haemoglobin subunits alpha and beta-1, potentially involved in physiological adaptations to deep water conditions. We downloaded data mapping to scaffolds 12, 81, and 215 (where the candidate genes are located) from the Cichlid Diversity Sequencing project (SRA accession: PRJEB1254). Variant calling was carried out as in ^21^. We used these data to examine a number of candidate genes highlighted by our analyses in more detail with respect to putatively functionally relevant nucleotide polymorphisms in *Diplotaxodon* as well as the greater Lake Malawi cichlid flock.

## Reproducibility statement

To ensure reproducibility of our analyses we have deposited detailed descriptions of the bioinformatics steps, custom scripts and further supplementary files (e.g. detailed morphological measurements, functional gene annotation results, GO enrichment tables) in a Github repository (https://github.com/HullUni-bioinformatics/Diplotaxodon_twilight_RAD; DOI: 10.5281/zenodo.259371). The raw RAD sequencing data is deposited with NCBI (Bioproject: PRJNA347810; SRA accessions: SRX2269491-SRX2269504). Whole genome resequencing data for the greater Lake Malawi flock is deposited on Genbank as part of the Cichlid diversity sequencing project (Bioproject accession: PRJEB1254).

## Author contributions

The paper was written by all authors. CH, MJG and DAJ conceived it, MJG collected and shared samples. Data analysis was carried out by CH.

## Acknowledgements

The RAD data was produced by Edinburgh Genomics, as a result of a NERC funded grant awarded to DAJ (NE/K000829/1). We would like to extend our thanks to Eric Miska, Richard Durbin and Milan Malinsky for permission to use the whole-genome data, assistance with variant calling, and comments on the manuscript.

## Supplementary

**Figure S1.**
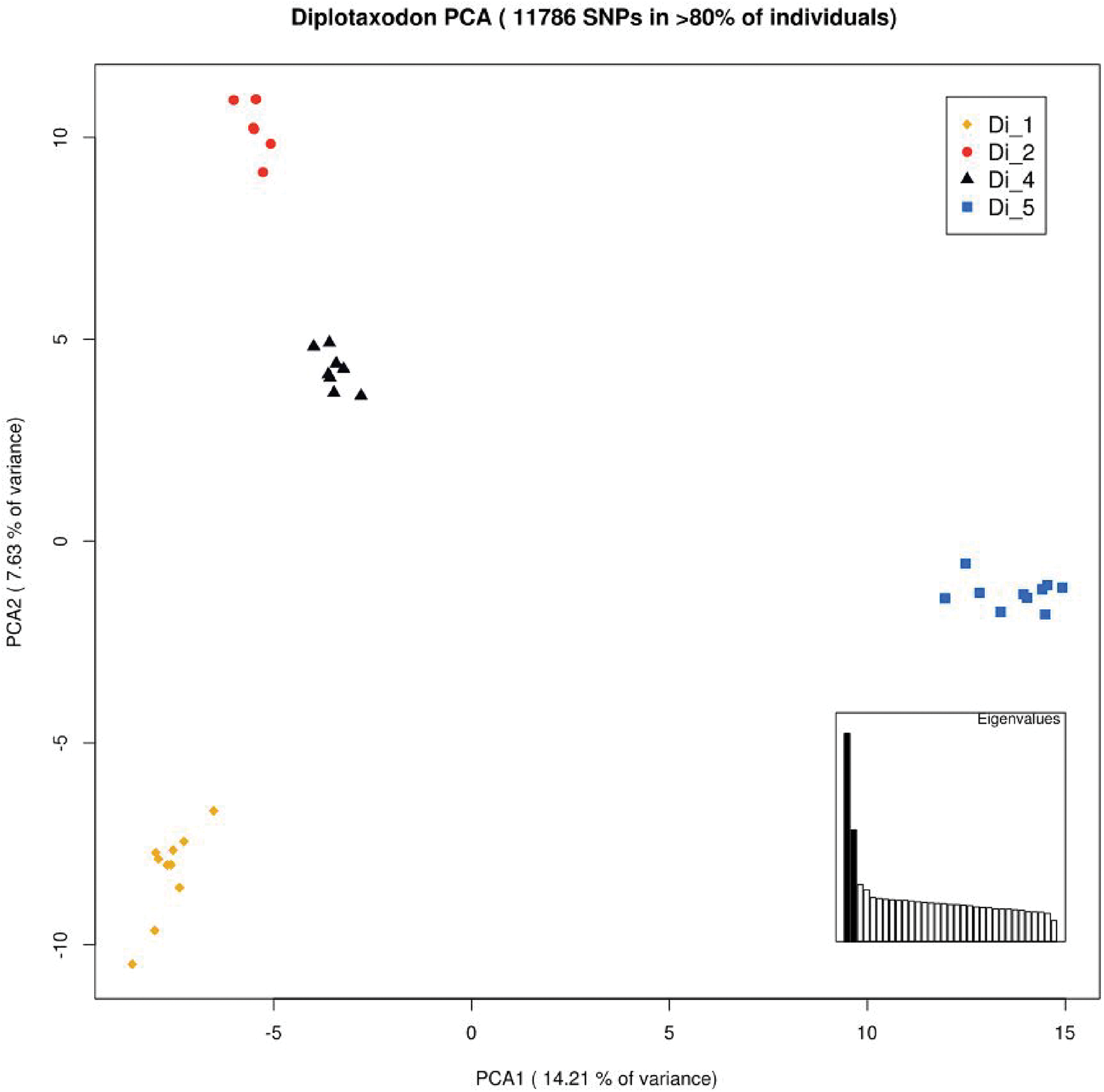
Principal component analysis based on 11,786 SNPs, *Diplotaxodon* species. Yellow - *D*. ‘macrops black dorsal’; red - *D*. ‘limnothrissa black pelvic’; black - *D*. ‘macrops offshore’; blue - *D*. ‘macrops ngulube’.

**Figure S2.**
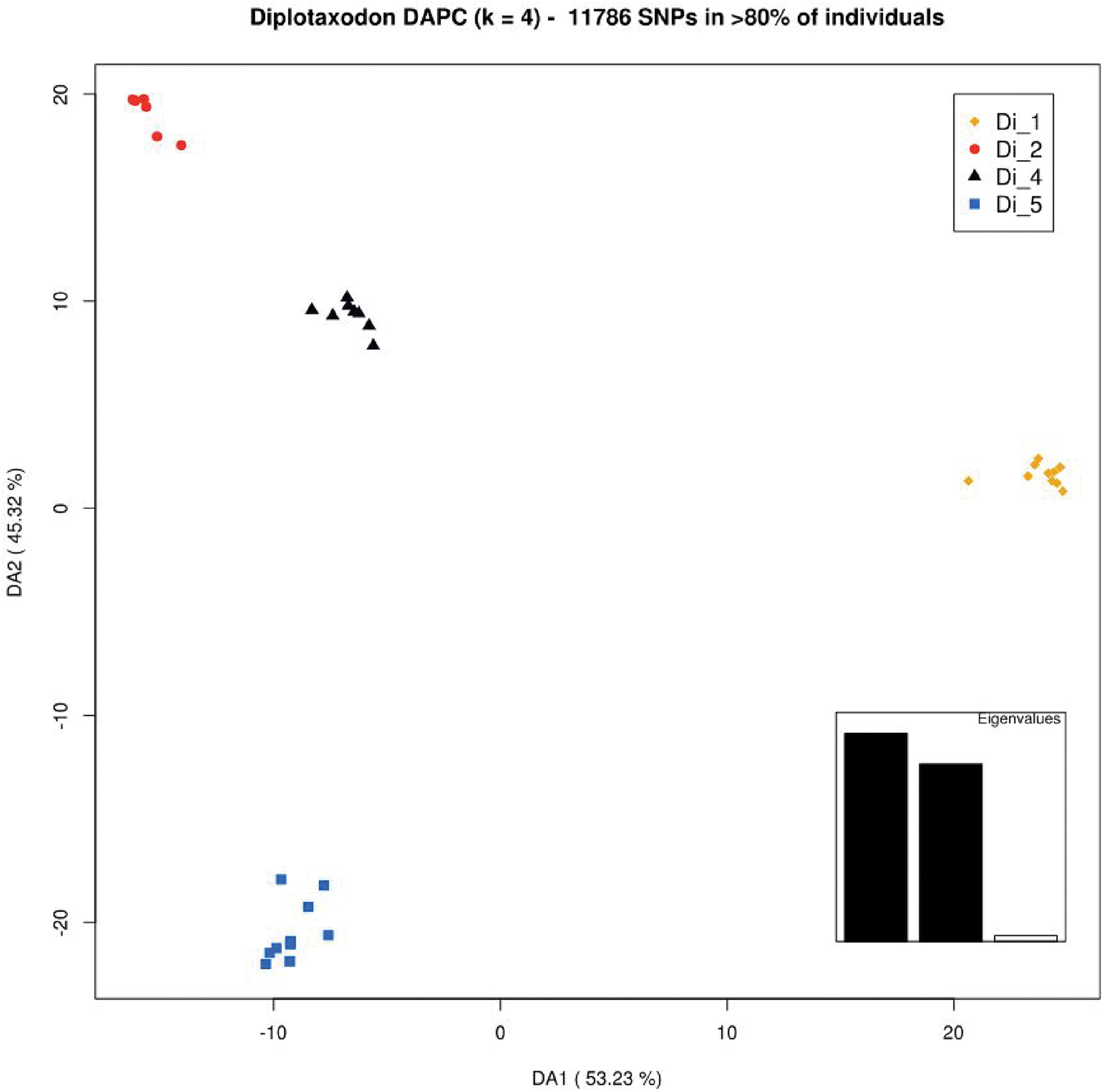
DAPC based on 11,786 SNPs, *Diplotaxodon* species. Yellow - *D*. ‘macrops black dorsal’; red - *D*. ‘limnothrissa black pelvic’; black - *D*. ‘macrops offshore’; blue - *D*. ‘macrops ngulube’.

**Figure S3.**
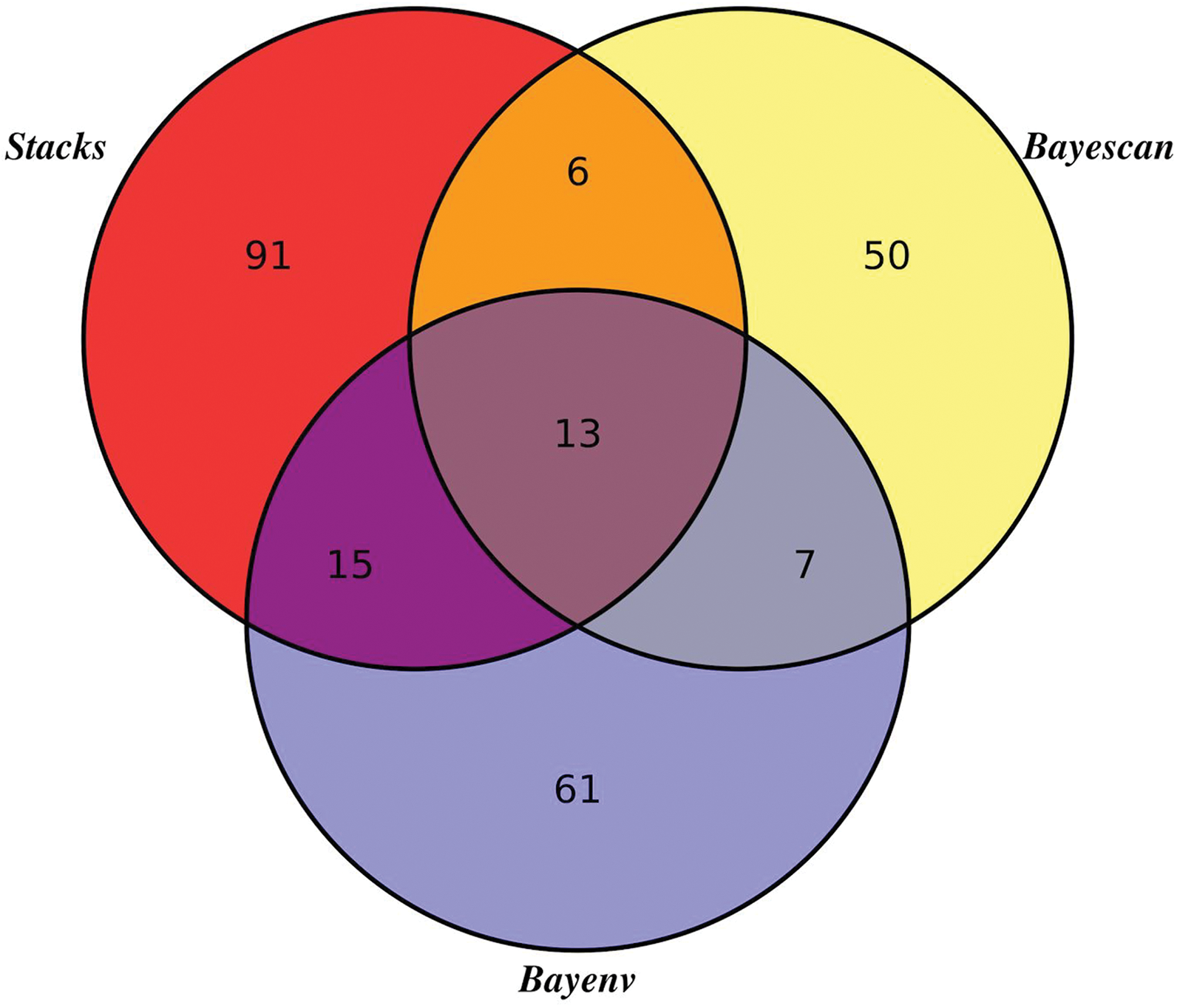
Number and concordance of candidate outlier loci highlighted by three independent approaches.

**Figure S4.**
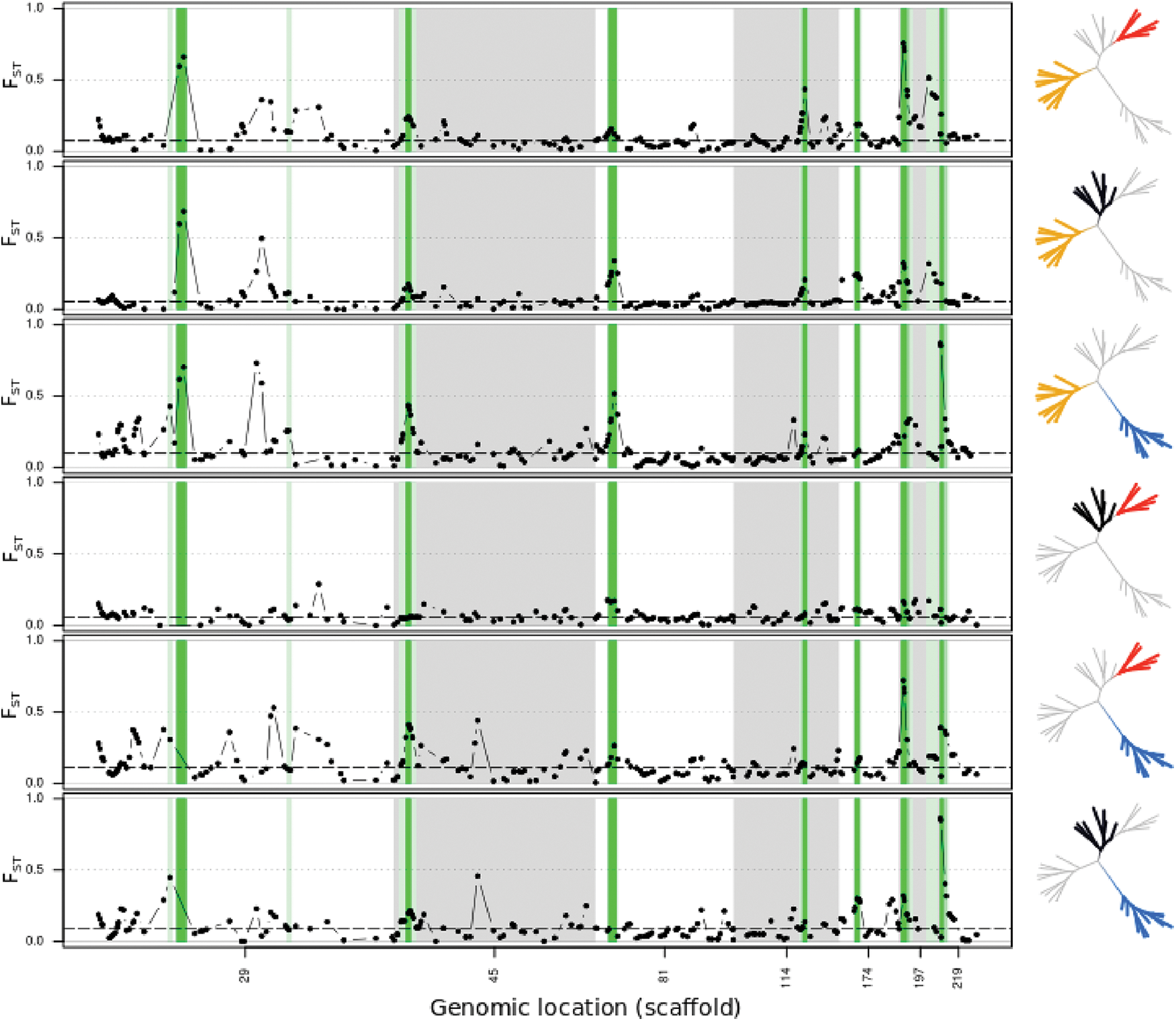
Pairwise Fst divergence at the six scaffolds (scaffold id on the x axis) containing loci highlighted as outliers by three independent detection approaches. Displayed are only scaffolds containing a minimum of 5 SNPs. Putative candidate regions are highlighted in shades of green. Dark green regions indicate support by three approaches (see tables 1 and S3 for summary of gene complements in highlighted regions). Dots represent the kernel smoothed averages across 50kb windows. Dashed lines indicate the genome wide Fst average. Population pairs are indicated by the highlighted regions in the phylogenetic trees on the right hand side, *Diplotaxodon* species. Yellow - *D*. ‘macrops black dorsal’; red - *D*. ‘limnothrissa black pelvic’; black - *D*. ‘macrops offshore’; blue - *D*. ‘macrops ngulube’.

**Figure S5.**
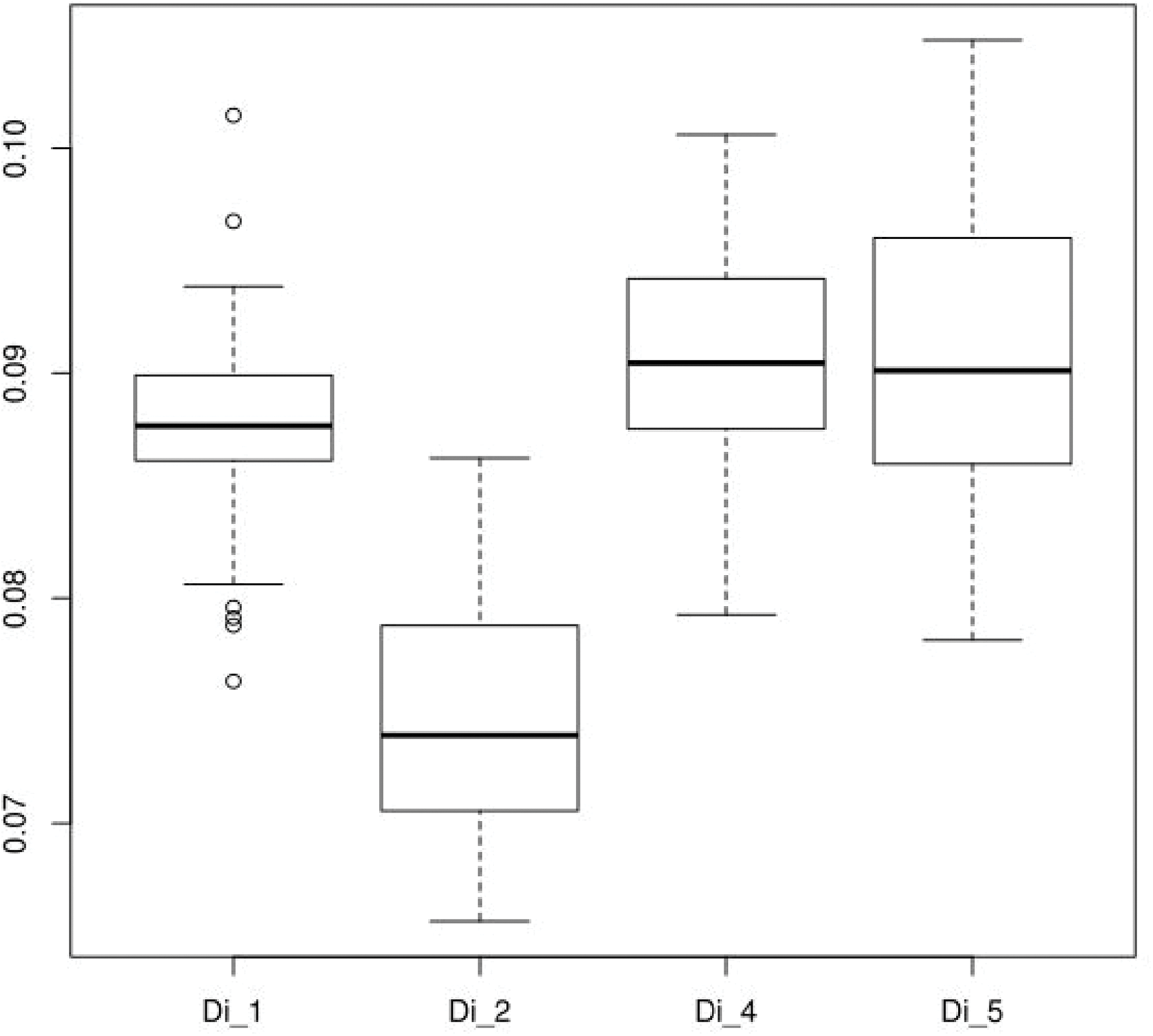
*Diplotaxodon* interspecific vertical eye diameter variation (normalized by total length of the fish).

